# Circadian Rhythms Time Seizure Severity in *Drosophila*

**DOI:** 10.1101/2025.06.11.659104

**Authors:** Mariam Huertas Radi, Adam A. Bradlaugh, Richard A. Baines

## Abstract

It is established that epilepsy patients can exhibit 24-hour rhythms in seizure severity and occurrence. While the pathways underlying seizure rhythmicity remain poorly understood, it seems likely that a contribution from the biological clock is involved. A better understanding of any such contribution may translate to better treatments. Here, the influence of the 24 h circadian rhythm on seizure activity in *Drosophila melanogaster* is investigated. Seizure-susceptible bang-sensitive *julius seizure* (*jus)* mutants were subjected to mechanically induced seizure at six different *zeitgeber* points. A clear sex-dependent phenotype was observed, with seizure severity showing a greater time-of-day effect in females, than males. The temporal pattern of seizure recovery time was bimodal, exhibiting both a morning and an evening peak. Rearing flies in constant light, which renders the molecular clock dysfunctional, abolished the seizure rhythm. Conversely, female seizure mutants reared in constant darkness, allowing free running of the circadian clock, continued to exhibit a bimodal rhythm of seizure severity. These findings support a role for the biological clock in seizure activity, at least in female *Drosophila*. Thus, this study validates *Drosophila* as a potential model for the identification of mechanisms modulating seizure rhythmicity, with the potential to aid future treatment of epilepsy.

## Introduction

Epilepsy is a prevalent neurological disorder, affecting ∼50 million people worldwide (WHO, 2024). Epilepsy is characterised by recurring seizures, the clinical manifestation of abnormal, excessive and/or synchronous neuronal activity in the brain (Fisher *et al*., 2014). The most common treatment is antiseizure medications (ASMs), the large majority of which target ion channels or neurotransmitter signalling, acting to restore excitatory-inhibitory balance (Löscher and Schmidt, 2011). However, despite more than thirty ASMs being developed, one third of people with epilepsy (PWE) remain drug refractory (Sultana *et al*., 2021). This underscores the need to identify novel targets and/or better ways to use existing ASMs.

Circadian rhythms are a fundamental aspect of biology; an internal timekeeping system that generates 24 h variations in most cellular, physiological and behavioural processes. Circadian rhythms drive temporal patterns in processes critical for the initiation and spread of seizures (Meisel *et al*., 2015). Thus, the relevance of circadian rhythms to epilepsy, and potential for treatment, is evident. Two of the most comprehensive databases of human seizures, SeizureTracker and NeuroVista, report 80% and 92% of PWE exhibiting seizures under circadian regulation, respectively (Karoly *et al*., 2018). Seizure occurrence and/or severity occur at fixed daily timings in most individuals. However, previous attempts at describing such temporal patterns have also uncovered significant variability (Leguia *et al*., 2021). Seizure rhythmicity is shown to depend on the cortical lobe in which seizure onset occurs, seizure type, age, chronotype of patients, and sleep-wake cycles (Kaleyias *et al*., 2011; Cho, 2012; van Campen *et al*., 2015). Altogether, these factors reduce the accuracy of circadian rhythm evaluation in mammals, as well as complicate the distinction of circadian rhythm effects on epilepsy from other influences.

The fruit fly, *Drosophila melanogaster*, provides a favourable model for studying circadian control of seizure severity (Johan Arief *et al*., 2018). Clock cells that orchestrate circadian rhythms in adult flies consist of ∼240 neurons (Reinhard *et al*., 2024), compared to the central circadian pacemaker suprachiasmatic nucleus with ∼100,000 neurons in humans (Swaab, 2003) and ∼20,000 neurons in mice (Varadarajan *et al*., 2018). Despite the evolutionary distance, the *Drosophila* and human genomes are highly conserved, and molecular pathways implicated in human diseases can be modelled and studied in *Drosophila*, including epileptiform activity (Cunliffe *et al*., 2015). For example, *bang senseless* (*bss*) *Drosophila* mutants, carrying a gain-of-function mutation in the *paralytic* (*para*) gene coding for a voltage-gated sodium channel (Na_v_), exhibit seizures that resemble those of Na_v_-associated pharmacologically resistant epilepsies in humans (Parker *et al*., 2011). More specifically, Zhang *et al*. (2018) reported that exposing *Drosophila* to the ASM phenytoin during nighttime decreased seizure burden compared with daytime administration, reproducing the nighttime application of phenytoin to PWE which similarly decreases seizure burden and associated adverse effects (Yegnanarayan, Mahesh and Sangle, 2006). Thus, the study of circadian rhythms contributing to seizure severity in fly seizure models has the potential to lead to significant progress in chronoepileptology.

Circadian rhythms are driven by clock genes, functioning as a molecular clock. Clock gene expression is altered in human epileptogenic brain tissue (Li *et al*., 2017; Wu *et al*., 2021), and the manipulation of clock genes influences the rhythmicity of seizures in mouse models (Gerstner *et al*., 2014). Therefore, clock genes have been suggested to be important determinants of seizure activity. Despite the differences between the mammalian and fly molecular clock components, forward genetic screens have revealed a conserved mechanism that operates in key central pacemaker neurons to generate circadian rhythms (Hardin, 2011; Stanton, Justin and Reitzel, 2022). Ever since the first clock gene *period (per)* was cloned and characterised in *Drosophila*, this organism has been a model of choice for understanding how the molecular clock regulates rhythmic neuronal activity. Overall, *Drosophila* provides significant potential for the study of clock genes and neuronal excitability, involved in seizure initiation and propagation.

In this study, we explored the interplay between circadian rhythms and seizure severity in *Drosophila*, to provide evidence of circadian rhythms contributing to seizure phenotypes in *Drosophila* models of epilepsy. We report that bang-sensitive (BS) *julius seizure* (*jus*) flies exhibit a bimodal circadian rhythm of seizure severity. This effect is larger and more pronounced in females. The influence of the circadian clock is abolished under constant light (LL), but not under constant darkness (DD) conditions, as would be predicted if seizure activity is regulated by the molecular clock. As a result, our findings spotlight *Drosophila* as a model for identifying the mechanism(s) modulating seizure rhythmicity.

## Materials and Methods

### Fly Stocks

Flies were maintained on a standard cornmeal medium (5 L water, 390 g glucose, 360 g maize, 250 g yeast, 40 g agar, 135 mL nipagin, 15 mL propionic acid) at 25°C under a 12:12 hour light-dark (LD) cycle, unless otherwise stated. Canton-Special (CS) wild type and w^-^; +; *jus*^*iso7*.*8*^ stocks were obtained from the Baines lab.

### Lighting Conditions

Adult flies (0-16 h post-eclosion) were collected using CO_2_, separated by sex and placed into new food vials in groups of six. Vials were either kept in a standard LD cycle or, when required, flies were moved to LL or DD. Seizure induction and locomotor assays were performed under LD, LL and DD. Light in LD was 1700 lux, and 1000 lux in LL, measured using a Thorlabs PM100A Power Meter.

### Seizure Induction: Vortex Assay

Seizure severity of 7 d old (± 16 h) flies (of both sexes) was measured via vortex assays (Kuebler and Tanouye, 2000). Testing was conducted at *zeitgeber* times (ZT) 0, 4, 8, 12, 16 and 20; ZT 0 indicating the onset of the light phase and ZT 12 the onset of the dark phase. In constant conditions, flies were tested at circadian times (CT). Flies were kept in LL or DD for 7 days prior to testing. Testing was conducted within one hour of each time point, equally spread across the 60 min for each lighting condition, genotype and sex. To assess seizure severity in mutants and CS (used as controls), food vials sealed with flugs (packed cotton bungs) were placed upside down (such that flies stood on the flugs) and vortexed on a standard laboratory vortex mixer at maximum speed for 10 s (Bermudez *et al*., 2023). A total of eight vials (six flies per vial), per ZT, were vortexed for each genotype and sex. Seizure recovery time was measured as time required to regain posture and mobility for each fly. Seizure recovery time was used as a proxy for seizure severity; a higher recovery time indicated a more severe seizure.

### Locomotor Activity and Sleep: DAM system

Circadian locomotor activity and sleep of female flies was measured using the Drosophila Activity Monitor (DAM) 5M software (TriKinetics Inc., Waltham, MA). Seizure mutant *jus* female flies (and CS used as controls) were collected upon eclosion (± 16 h), anesthetised with CO_2_, and transferred individually into DAM tubes (5 mm diameter, 65 mm length) using a fine paintbrush. Tubes were prepared by adding ∼5 mm standard cornmeal medium (see above) to one end of the tube, sealed with a rubber cap, and a small cotton plug to the other end. Before any recordings started, flies were allowed 24 h to adapt to the new tube. DAM monitors with 32 flies per genotype, and per lighting condition, were initially entrained for 3 days in LD, followed by 5 days of either DD or LL. Experiments were run at 25°C, and an environmental monitor (DEnM; Trikinetics Inc.) was used to monitor conditions. Locomotor activity of flies was summed into 1-min bins. Flies that died before the end of the experiment were excluded from analysis. In all experiments, data are from flies collected and run concurrently (per lighting condition). DAM circadian and sleep data were analysed using the Sleep and Circadian Analysis Matlab Program (SCAMP, MATLAB 24.2) (Vecsey *et al*., 2024).

### Statistical Analysis

Statistics were performed in GraphPad Prism (10.4.1). All data obtained were tested for normality and equal variances using Shapiro-Wilk and Brown-Forsythe tests, respectively, before any further statistical analyses were carried out. Sample sizes for vortex assays were determined from previous studies (Mituzaite *et al*., 2021). Outlier tests were performed on vortex assay data to remove obvious outliers following ROUT 1% criteria. To determine the effect of ZT or CT (in the case of LL and DD data) on seizure recovery time, one-way ANOVA (or the equivalent non-parametric Kruskal-Wallis test, or Welch’s ANOVA for unequal variances) were used. Cubic spline curves were generated from means to visualize circadian rhythms in seizure severity. To compare recovery times between lighting conditions, two-way ANOVA was performed. To compare circadian locomotor activity and sleep data obtained from SCAMP between lighting conditions, one-way ANOVA (or the equivalent non-parametric Kruskal-Wallis test, or Welch’s ANOVA for unequal variances) was performed. Flies were considered rhythmic when the calculated rhythmicity index (RI) was greater than 0.3, weakly rhythmic for RI values between 0.1 and 0.3, and arrhythmic when RI was less than 0.1 (Sundram *et al*., 2012). The level of significance accepted for all tests was *p* ≤ 0.05. For significant test results, the appropriate *post hoc* test was performed.

## Results

### Circadian Bimodal Pattern of Seizure Recovery in Females

The aim of this study was to establish the influence of circadian timing on seizure severity in *Drosophila*. To address this, BS seizure-susceptible *jus* mutants were used. These mutants are defective for an uncharacterized protein-coding gene *CG14509* (Horne *et al*., 2017) and show prolonged seizures when subjected to a strong mechanical shock that simultaneously activates sensory input from peripheral hairs. However, this mutation does not show spontaneous seizures. Male and female *jus* flies, under (12:12) LD conditions, were subjected to vortex assay to induce seizures at ZT 0, 4, 8, 12, 16 and 20, and recovery times measured (Fig. 1A). The recovery time of CS, measured at the same time, was significantly lower than *jus* mutants, independent of sex (Kruskal-Wallis tests, *p* < 0.0001), thus validating the seizure phenotype observed.

**Figure 1.**
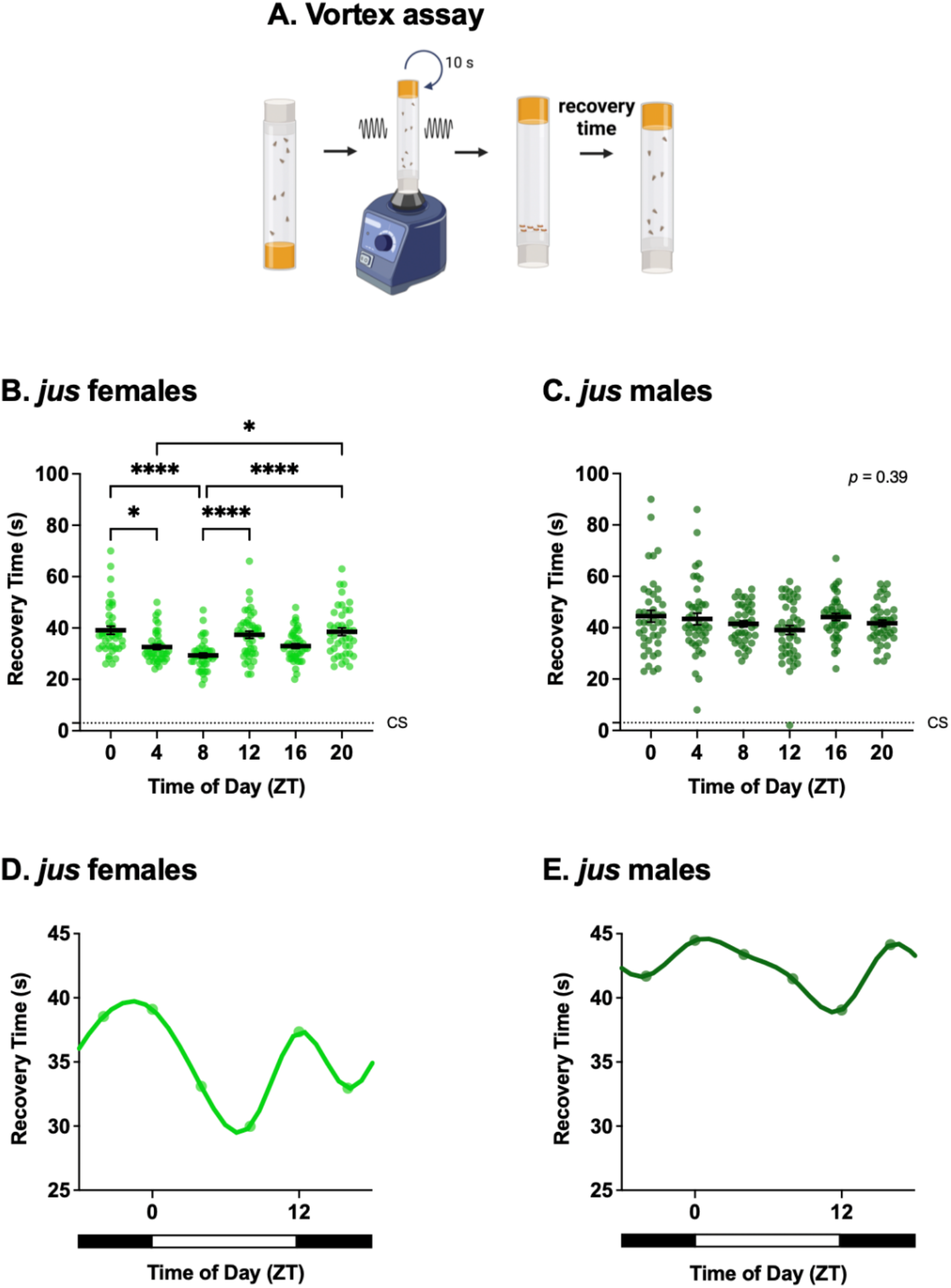
Effect of *zeitgeber* time (ZT) on seizure recovery time of *jus* seizure mutants. Recovery time (s) of adult flies in 12:12 light-dark conditions, tested via the vortex assay at ZT 0, 4, 8, 12, 16 and 20. (**A**) Schematic of the vortex assay. Created in BioRender. Huertas Radi, M. (2025) https://BioRender.com/u6v9tc9. (**B**) *jus* females show a statistically significant effect of ZT on seizure recovery time (Kruskal-Wallis test, *H (5) = 43*.*92, p < 0*.*0001, n = 39-43*). (**C**) *jus* males show no statistically significant effect of ZT on recovery time (Kruskal-Wallis test, *H (5) = 5*.*23, p = 0*.*39, n = 41-43*). For B-C, individual dots are recovery times of individual flies vortexed only once, derived from a total of eight vials (six flies per vial) per ZT and per sex. Average seizure recovery time of wild type CS controls are shown by dotted lines and are significantly different to *jus* female and male mutants (Kruskal-Wallis test, *H (6) = 109*.*0* (females), *84*.*65* (males), *p < 0*.*0001*, for both). Data are presented as mean ± SEM. Statistically significant comparisons from Dunn’s multiple comparison tests are indicated as ^****^ *p* ≤ 0.0001, ^***^ *p* ≤ 0.001, ^**^ *p* ≤ 0.01, ^*^ *p* ≤ 0.05. Full descriptive statistics and *p* values are reported in Table 1. (**D**) Circadian cycle of seizure severity of female *jus* exhibits a bimodal circadian rhythm. (**E**) Circadian cycle of seizure severity of male *jus* exhibits a similar phase to females but with a smaller effect size. For D-E, data are presented as means of B-C vortex data for each ZT. Horizontal white and black bars indicate when lights were on (ZT 0 to ZT 12) and off (ZT 12 to ZT 24).

Recovery time from induced seizure in *jus* showed a sex-dependent bias. Here, *jus* females showed a statistically significant effect of ZT on seizure recovery (Fig. 1B, Kruskal-Wallis test, *p* < 0.0001). Dunn’s multiple comparison tests revealed significant differences between ZT 0-4, ZT 0-8, ZT 4-20, ZT 8-12, and ZT 8-20 (full descriptive statistics and *p* values are reported in Table 1). By contrast, males showed no statistically significant differences in recovery time (i.e., seizure severity) across time points (Fig. 1C, Kruskal-Wallis test, *p* = 0.39). To further examine the temporal pattern of *jus* seizure severity, spline curves were generated from vortex assay data. In females, cycles of seizure recovery exhibited a bimodal rhythm, exhibiting two peaks (ZT = 12.5 h and ZT = 22.7 h) and two troughs (ZT = 6.9 h and ZT = 16.2 h) across the 24 h day (Fig. 1D). Notably, spline curves generated for males also showed a similar temporal pattern, however, with a much smaller effect size (Fig. 1E). Therefore, we conclude that seizure severity tested by the vortex assay is influenced by ZT, with an effect size that is larger in female *jus* mutants.

**Table 1.** Full descriptive statistics of multiple comparison tests of seizure severity between timepoints.

| Lighting Condition | ZT/CT | Mean recovery time (s) | SEM | post hoc test | *p* value | Significance | Fig. |
| --- | --- | --- | --- | --- | --- | --- | --- |
| LD | 0 | 39.10 | 1.57 | Dunn's | < 0.05 | * | 1B |
|  | 4 | 32.58 | 0.96 |  |  |  |  |
|  | 0 | 39.10 | 1.57 | Dunn's | < 0.0001 | **** | 1B |
|  | 8 | 29.33 | 0.94 |  |  |  |  |
|  | 4 | 32.58 | 0.96 | Dunn's | < 0.05 | * | 1B |
|  | 20 | 38.54 | 1.48 |  |  |  |  |
|  | 8 | 29.33 | 0.94 | Dunn's | < 0.0001 | **** | 1B |
|  | 12 | 37.35 | 1.35 |  |  |  |  |
|  | 8 | 29.33 | 0.94 | Dunn's | < 0.0001 | **** | 1B |
|  | 20 | 38.54 | 1.48 |  |  |  |  |
| DD | 4 | 64.89 | 4.62 | Dunnett's | < 0.05 | * | 2C |
|  | 8 | 48.95 | 2.32 |  |  |  |  |
|  | 4 | 64.89 | 4.62 | Dunnett's | < 0.05 | * | 2C |
|  | 20 | 48.35 | 2.24 |  |  |  |  |
|  | 16 | 62.08 | 3.71 | Dunnett's | < 0.05 | * | 2C |
|  | 20 | 48.35 | 2.24 |  |  |  |  |

### Bimodal Rhythm Abolished in Constant Light but Maintained in Constant Darkness

To provide additional evidence for a contribution of circadian time to seizure recovery, we determined the effects of altering the molecular clock. Under conditions of constant light (LL), *Drosophila* are arrhythmic because of continuous degradation of the protein TIMELESS (TIM) by its interaction with CRYPTOCHROME (CRY) (Marrus, Zeng and Rosbash, 1996). Thus, the expectation, in keeping with a contribution from the molecular clock, is that under LL a bimodal pattern of seizure severity would be absent. We focused attention to *jus* females because of the larger effect size. Indeed, *jus* females reared under LL showed no significant difference in recovery times between circadian timepoints (CT) (Fig. 2A-B, Kruskal-Wallis test, *p* = 0.26). Therefore, time of the day did not influence seizure severity of *jus* females in LL.

**Figure 2.**
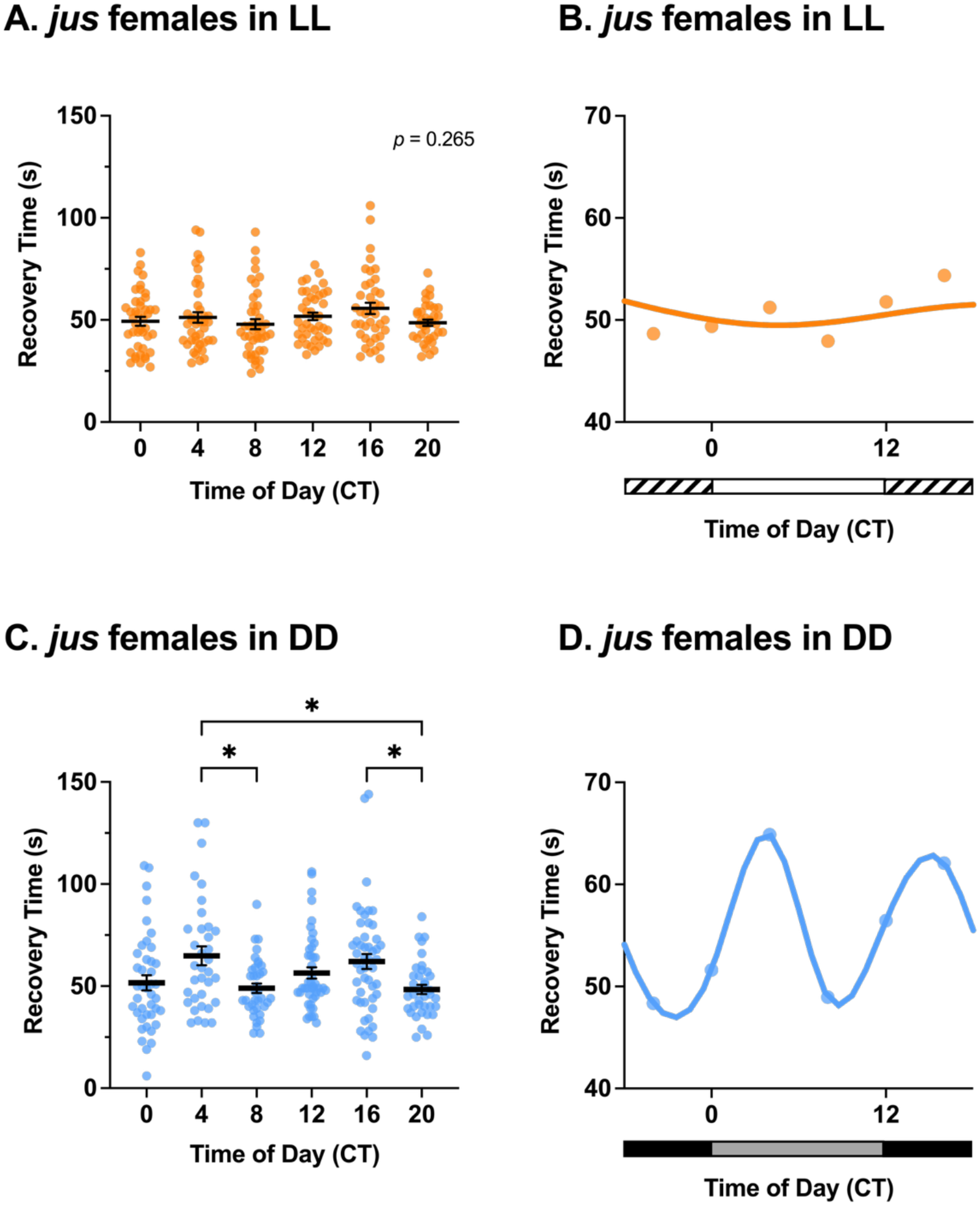
Rhythmicity of *jus* female seizure severity is abolished in constant light (LL) but maintained in constant darkness (DD). (**A**) Seizure recovery times (s) of *jus* females tested via the vortex assay after 7 days of LL show no effect of circadian time (CT) (Kruskal-Wallis test, *H (5) = 6*.*45, p = 0*.*26, n = 37-43*). (**B**) Temporal pattern of seizure severity of *jus* females, reared in LL, shows no obvious rhythmicity. (**C**) *jus* females under DD show a significant effect of circadian time on seizure recovery time (Welch’s ANOVA, *W (5, 108*.*92) = 4*.*23, p < 0*.*01, n = 35-49*). Statistically significant comparisons from Dunnett’s multiple comparison tests are indicated as * *p* ≤ 0.05, full descriptive statistics and *p* values are reported in Table 1. (**D**) Circadian cycle of seizure severity of *jus* females reared in DD exhibits a bimodal circadian rhythm. For A and C, individual dots are recovery times of individual flies vortexed only once, derived from a total of eight vials (six flies per vial), per CT and per lighting condition. Data are presented as mean ± SEM. For B and D, data are presented as means of A and C, respectively. In B, horizontal white and stripped bars represent periods of light and ‘subjective nighttime’ in LL, respectively. In D, horizontal black and grey bars represent periods of dark and ‘subjective daytime’ in DD, respectively.

Under conditions of constant darkness (DD), the *Drosophila* molecular clock free-runs (Konopka and Benzer, 1971). Maintenance of a bimodal rhythm of seizure severity under DD would, thus, further support our hypothesis of a contribution from the molecular clock to seizure severity. Indeed, seizure recovery times of *jus* females maintained under DD were significantly influenced by CT (Fig. 2C-D, Welch’s ANOVA, *p* < 0.01). Dunnett’s multiple comparison tests showed significant differences between recovery times at CT 4-8, CT 4-20, and CT 16-20 (full descriptive statistics and *p* values are reported in Table 1). Therefore, time of the day influenced seizure severity of *jus* females in DD. The temporal pattern of seizure severity was circadian, exhibiting two peaks (CT = 4.1 h and CT = 15.3 h) and two troughs (CT = 8.7 h and CT = 21.8 h) across the 24 h day (Fig. 2D). However, the timing of peaks and troughs, and therefore the phase of the DD cycle, was shifted to ∼4 h later relative to the LD cycle. Taken together, these results validate the hypothesis that the circadian clock governs seizure severity in *jus* females.

### Circadian Locomotor Activity Complements Seizure Rhythmicity Findings

To further support the possible contribution of the circadian clock to seizure activity, locomotor activity of *jus* females was measured via the TriKinetics system under the same lighting conditions as vortex assays (Fig. 3A-C). In LD conditions, adult flies showed an expected bimodal circadian rhythm of locomotion, exhibiting morning and evening peaks of activity at ZT 0 and ZT 12 (Fig. 3A). The average period length was 23.95 h, and 89.3% of flies were rhythmic (88% of which were strongly rhythmic), with an average 0.36 ± 0.03 rhythmicity index (RI) and 2.82 ± 0.21 rhythmicity strength (RS). Conversely, when kept in LL conditions, 48% of *jus* females were arrhythmic and 52% were only weakly rhythmic (Fig. 3B). Locomotor activity of *jus* females in LL was below standard rhythmicity thresholds, with a 0.091 ± 0.02 RI and 0.72 ± 0.13 RS. Lastly, when kept under free-running DD conditions, 96.3% of flies were rhythmic (76.9% of which were strongly rhythmic) (Fig. 3C). The average period length of flies was 24.19 h, with an average 0.33 ± 0.02 RI and 2.61 ± 0.15 RS. Here, it is worth noting that the activity profile of *Drosophila* in DD does not exhibit two distinctive peaks of morning and evening activity as in LD, but rather one merged peak of activity.

**Figure 3.**
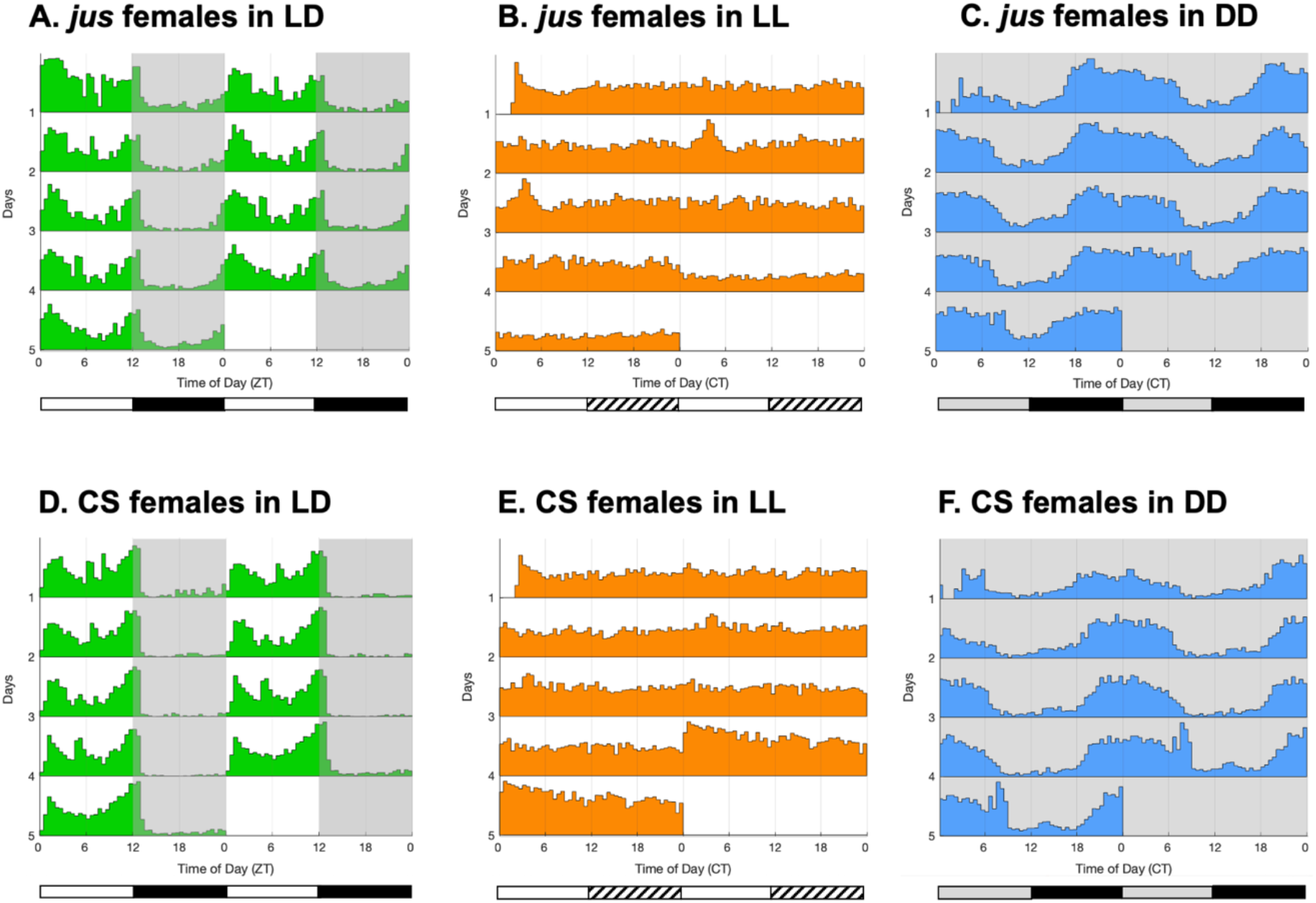
Double-plotted actograms of circadian locomotor activity of *jus* and CS females in 12:12 light-dark (LD), constant light (LL), and constant darkness (DD) conditions. Activity of adult flies was analysed via the TriKinetics system. (**A**) Locomotor activity of *jus* females in 12:12 LD shows a bimodal circadian rhythm, with peaks of activity at ZT 0 and ZT 12 (n = 28). (**B**) Locomotor activity of *jus* females in LL (switched to LL after 3 days of LD) shows arrhythmicity, with a constant level of activity lacking any temporal pattern (n = 25). (**C**) Locomotor activity of *jus* females in DD (switched to DD after 3 days of LD) shows a circadian rhythm, but not bimodal due to the lack of a ‘siesta’ period (n = 27). (**D**) Locomotor activity of CS females in 12:12 LD shows a bimodal circadian rhythm, with peaks of activity at ZT 0 and ZT 12 (n = 16). (**E**) Locomotor activity of CS females in LL (switched to LL after 3 days of LD) shows arrhythmicity, with a constant level of activity lacking any temporal pattern (n = 26). (**F**) Locomotor activity of CS females in DD (switched to DD after 3 days of LD) shows a circadian rhythm, but not bimodal due to the lack of a ‘siesta’ period (n = 24). In actograms, lights off are indicated by grey background shading, and lights on are indicated by white shading. Horizontal white and black bars under LD actograms indicate when the lights were on (ZT 0 to ZT 12) and off (ZT 12 to ZT 24). Horizontal white and stripped bars under LL actograms represent periods of light and ‘subjective nighttime’ in LL, respectively. Horizontal black and grey bars under DD actograms represent periods of dark and ‘subjective daytime’ in DD, respectively.

As controls, CS female flies were monitored under the same conditions (Fig. 3D-F). In LD conditions, adult flies showed a circadian rhythm, like *jus* females, exhibiting morning and evening peaks of activity at ZT 0 and ZT 12 (Fig. 3D). The average period length was 24.03 h, and 100% of flies were rhythmic (93.8% of which were strongly rhythmic), with an average 0.44 ± 0.02 RI and 3.43 ± 0.17 RS. In LL conditions, 69.2% were arrhythmic and 30.8% were only weakly rhythmic (Fig. 3E). Locomotor activity of CS females in LL was below standard rhythmicity thresholds, with a 0.083 ± 0.02 RI and 0.65 ± 0.19 RS. On the contrary, when kept under free-running DD conditions, 95.8% of flies were rhythmic (69.6% of which were strongly rhythmic), the average period length was 24.75 h, and a 0.32 ± 0.02 RI and 2.51 ± 0.18 RS. (Fig. 3F). Thus, *jus* females described above behave as control CS flies. Here, it is worth noting that fly lines used for activity monitoring were not isogenic lines, thus meaningful comparisons can only be made within genotypes between conditions.

Comparisons of rhythmicity values of *jus* females across lighting conditions revealed RI and RS values in LL significantly lower than in LD and DD (Kruskal-Wallis with Dunn’s multiple comparison tests, *p* < 0.0001 for both). No significant differences were found between rhythmicity values in LD and DD (Kruskal-Wallis with Dunn’s multiple comparison test, *p* = 0.85). Therefore, this pattern of circadian locomotor activity validates that, as expected, the behaviour of *jus* females in LD and DD conditions is rhythmic, with a period of around 24 h, whilst arrhythmic in LL. Thus, complementing the loss of seizure severity circadian rhythms when *jus* females were reared in LL prior to vortex testing.

### LL and DD Increase Seizure Severity and Involve Sleep Disturbances

To determine whether LL and DD conditions had varying effects on the seizure severity of *jus* females, comparisons between these two lighting conditions to LD conditions were performed. LL and DD showed an increase in seizure severity (e.g., a longer recovery time) in *jus* females compared to LD, independent of circadian time (Fig. 4A, two-way ANOVA, *p* < 0.0001). Dunnett’s multiple comparison tests showed a significant increase in seizure severity in LL and DD at all timepoints tested (full descriptive statistics and *p* values are reported in Table 2). Thus, both altered light regimes increase seizure severity of *jus* females, in addition to effects related to circadian rhythmicity.

**Table 2.** Full descriptive statistics of multiple comparison tests of seizure severity between LL and DD to control LD conditions.

| ZT/CT | Lighting Condition | Mean recovery time (s) | SEM | post hoc test | *p* value | Significance | Fig. |
| --- | --- | --- | --- | --- | --- | --- | --- |
| **0** | LD | 39.10 | 1.57 |  |  |  | 4A |
|  | LL | 49.37 | 2.17 | Dunnett's | < 0.01 | ** |  |
|  | DD | 51.61 | 3.69 | Dunnett's | < 0.001 | *** |  |
| **4** | LD | 32.58 | 0.96 |  |  |  | 4A |
|  | LL | 51.24 | 2.59 | Dunnett's | < 0.0001 | **** |  |
|  | DD | 64.89 | 4.62 | Dunnett's | < 0.0001 | **** |  |
| **8** | LD | 29.33 | 0.94 |  |  |  | 4A |
|  | LL | 47.93 | 2.47 | Dunnett's | < 0.0001 | **** |  |
|  | DD | 48.95 | 2.32 | Dunnett's | < 0.0001 | **** |  |
| **12** | LD | 37.35 | 1.35 |  |  |  | 4A |
|  | LL | 51.78 | 1.85 | Dunnett's | < 0.0001 | **** |  |
|  | DD | 56.44 | 2.76 | Dunnett's | < 0.0001 | **** |  |
| **16** | LD | 32.95 | 0.90 |  |  |  | 4A |
|  | LL | 55.65 | 2.78 | Dunnett's | < 0.0001 | **** |  |
|  | DD | 64.1 | 4.16 | Dunnett's | < 0.0001 | **** |  |
| **20** | LD | 38.54 | 1.48 |  |  |  | 4A |
|  | LL | 48.65 | 1.54 | Dunnett's | < 0.05 | * |  |
|  | DD | 48.35 | 2.24 | Dunnett's | < 0.05 | * |  |

**Figure 4.**
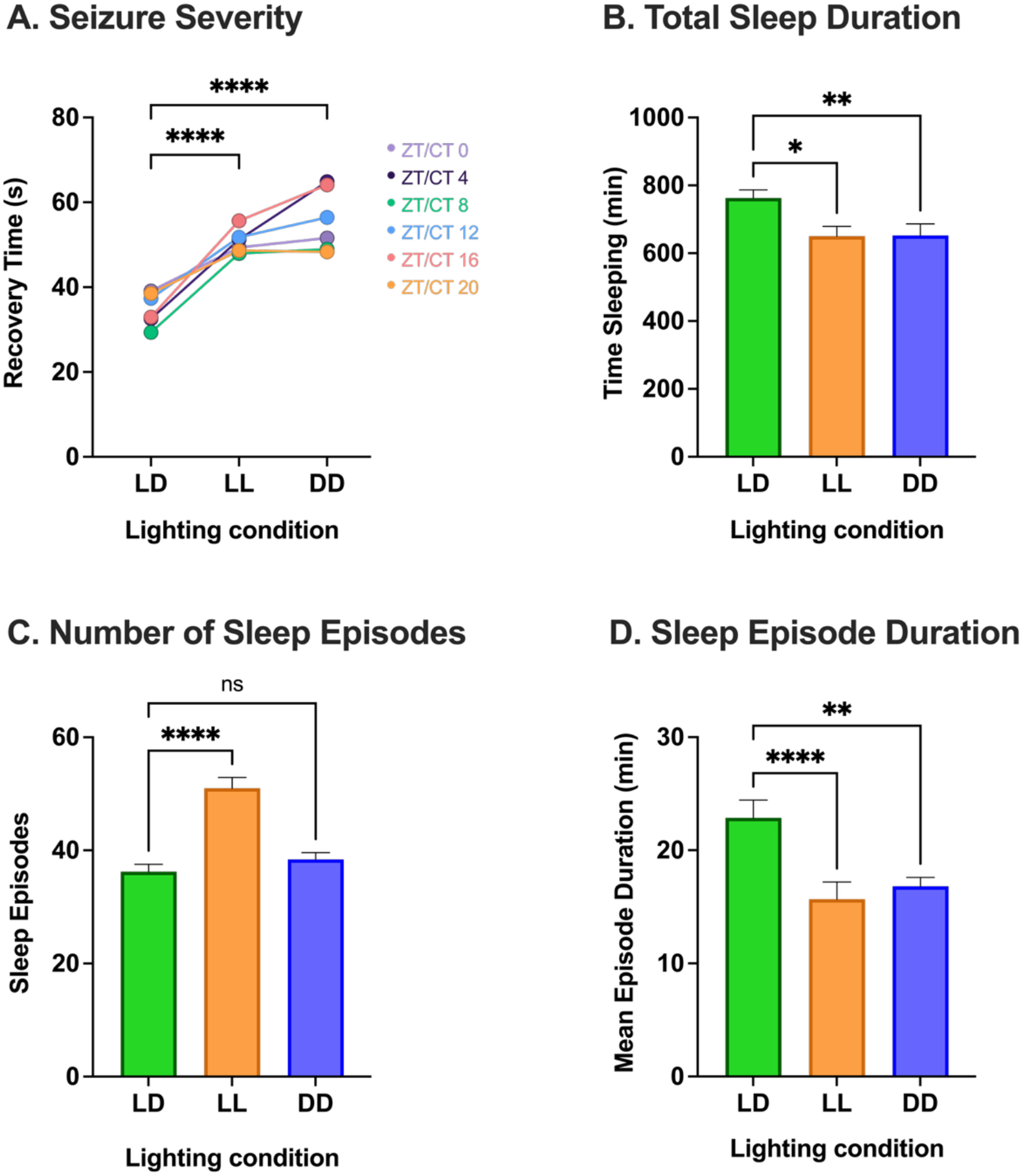
Constant light (LL) and constant darkness (DD) show an increase in seizure severity in *jus* females, potentially linked to sleep loss and fragmentation. (**A**) Lighting condition has a statistically significant effect on seizure severity of *jus* females tested via the vortex assay. Seizure recovery times in LL and DD show an increase in seizure severity compared to 12:12 LD conditions at all timepoints (two-way ANOVA, *F (2, 719) = 111*.*0, p < 0*.*0001, n = 6 timepoints*). Full descriptive statistics and *p* values of multiple comparison tests are reported in Table 2. Each dot represents the mean seizure recovery time for each ZT/CT, per lighting condition. Individual timepoints are coloured differently and joined across lighting conditions with lines. LD vortex data is taken from Fig. 1B, LL from Fig. 2A, and DD from Fig. 2C. (**B**) Total sleep duration of *jus* females reared in LL (in orange) and DD (in blue) was significantly lower than in LD (in green) (Kruskal-Wallis test, *H (2) = 11*.*87, p < 0*.*01, n = 25-28*). (**C**) Number of sleep episodes of *jus* females monitored in LL and DD was significantly higher than in LD (one-way ANOVA, *F = 28*.*50, p < 0*.*0001, n = 25-28*). (**D**) Duration of sleep episodes of *jus* females kept in LL and DD were significantly lower than in LD (Kruskal-Wallis test, *H (2) = 28*.*05, p < 0*.*0001, n = 25-28*). Statistically significant comparisons from Dunnett and Dunn’s multiple comparison tests are indicated as ^****^ *p* ≤ 0.0001, ^***^ *p* ≤ 0.001, ^**^ *p* ≤ 0.01, ^*^ *p* ≤ 0.05. Full descriptive statistics and *p* values of multiple comparison tests are reported in Table 3. Data are presented as mean ± SEM.

Potentially, an increase in seizure severity could be related to sleep loss of *jus* females when kept in LL and DD conditions. Preclinical and clinical studies have shown that sleep deprivation can increase the risk of epileptic seizures in humans (Bonilla-Jaime *et al*., 2021; Dell’Aquila and Soti, 2022), hence a similar mechanism could explain the increase in seizure burden observed in *jus* females. Sleep characteristics of *jus* females were monitored via the TriKinetics system during 5 days under LD, LL and DD lighting conditions. Here, it is worth noting that fly lines used for sleep monitoring were not isogenic. However, the following comparisons were performed within the same genotype, between different conditions. On average, the total amount of sleep *jus* females had per day in LL and DD was significantly lower than in LD (Fig. 4B, Kruskal-Wallis with Dunn’s multiple comparison test, *p* < 0.05 and *p* < 0.01, respectively, full descriptive statistics and *p* values are reported in Table 3). Regarding the amount of sleep episodes, *jus* females underwent a significantly higher number of sleep bouts per day in LL (Fig. 4C, one-way ANOVA with Dunnett’s multiple comparison test, *p* < 0.0001), but not in DD (Fig. 4C, one-way ANOVA with Dunnett’s multiple comparison test, *p* = 0.47, full descriptive statistics and *p* values are reported in Table 3). In terms of mean duration of sleep episodes, flies in LL and DD took significantly shorter sleep bouts than in LD (Fig. 4D, Kruskal-Wallis test with Dunn’s multiple comparison test, *p* < 0.0001 and *p* < 0.01, respectively, full descriptive statistics and *p* values are reported in Table 3). Therefore, *jus* females in LL and DD exhibited more fragmented sleep, i.e., a higher number, but shorter duration, of sleep bouts, as well as sleep loss as indicated by the reduced total amount of sleep. Hence, such sleep disturbances could exacerbate seizure severity of *jus* females when tested via the vortex assay.

**Table 3.** Full descriptive statistics of multiple comparison tests of sleep parameters between LL and DD to control LD conditions.

| Sleep parameter | Lighting Condition | Mean | SEM | post hoc test | *p* value | Significance | Fig. |
| --- | --- | --- | --- | --- | --- | --- | --- |
| **Total Sleep Duration (min)** | LD | 763.0 | 23.77 |  |  |  | 4B |
|  | LL | 650.2 | 28.79 | Dunn's | < 0.05 | * |  |
|  | DD | 652.2 | 34.52 | Dunn's | < 0.01 | ** |  |
| **Number of Sleep Episodes** | LD | 36.24 | 1.30 |  |  |  | 4C |
|  | LL | 51.0 | 1.90 | Dunn's | < 0.0001 | **** |  |
|  | DD | 38.42 | 1.19 | Dunn's | = 0.47 | ns |  |
| **Sleep Episode Duration (min)** | LD | 22.87 | 1.58 |  |  |  | 4D |
|  | LL | 15.67 | 0.80 | Dunn's | < 0.0001 | **** |  |
|  | DD | 16.81 | 1.53 | Dunn's | < 0.01 | ** |  |

## Discussion

The aim of this study was to determine whether circadian timing has an effect on *Drosophila* seizure severity. To achieve this, recovery times of *jus* mutants were recorded at different times of the day. Male flies showed no effect of time on seizure severity, whereas females revealed an influence of circadian time on seizure severity measured via the vortex assay. In humans, sex differences are evident in many epilepsies and seizure conditions (Hophing, Kyriakopoulos and Bui, 2022). Neurotransmission and synaptic function, which mediate neural activity, and thus seizure activity, are modulated in a circadian-dependent and sex-specific manner (Logan *et al*., 2022). Current evidence suggests that males exhibit greater overall susceptibility to excitability episodes and occurrence of seizures, whilst females exhibit greater fluctuations in seizure occurrence (Reddy, Thompson and Calderara, 2021). The precise molecular mechanisms of this remain unknown, although most studies suggest the involvement of neuroendocrine factors (Wang, Zhuo and Wang, 2023). The only sex differences reported in *Drosophila* relating to epilepsy are at the transcriptomic level in pentylenetetrazol-induced seizure models, where females showed downregulation of genes related to the ribosomal pathway (Sharma, Mohammad and Singh, 2009). Thus, behavioural differences between males and females in seizure rhythmicity have not been previously reported. Given the sex differences observed in this study, future work on *Drosophila* male and female hormones could perhaps offer an opportunity to elucidate the mechanisms underlying sex differences in epilepsy.

Seizure severity of *jus* females followed a bimodal circadian rhythm, with two peaks and two troughs across the 24 h. In the context of *Drosophila*, the daily locomotor activity rhythm also follows a bimodal circadian rhythm, as shown by the circadian locomotor activity results of this study. Activity peaks at lights-on, ZT 0 termed “morning peak”, and at lights-off, ZT 12 termed “evening peak” (Miyasako et *al*., 2007; Chiu et *al*., 2010), precisely match the seizure severity peaks observed in *jus* female mutants at ZT 12, and close to ZT 24 (= ZT 0). Pigment Dispersing Factor (PDF)-positive small ventrolateral neurons (s-LNvs) are the main pacemakers for *Drosophila* circadian activity. Interestingly, these cells play a major role in driving morning peaks of activity (Miyasako, Umezaki and Tomioka, 2007), as well as in the regulation of the phase of other cells, such as pre-motor neurons (Liang *et al*., 2019). Peptides released by central pacemaker neurons (e.g., DH44 and Hugin) link the clock to motor outputs (King *et al*., 2017), thus modulating the locomotor activity rhythm. Increased synaptic excitation of motoneurons is associated with seizure activity in *jus* mutants (Marley and Baines, 2011). Hence, potentially, increased PDF-positive s-LNvs neural activity and excitability could underlie seizure peaks: thus, seizure burden following a bimodal circadian rhythm similar to locomotor activity. In addition, the vortex assay induces seizures by overstimulating *Drosophila* sensory bristles (Kuebler and Tanouye, 2000). Circadian regulation of sensory inputs, such as circadian rhythms in light sensitivity of *Drosophila* (Vinayak *et al*., 2013), could be involved in driving the seizure severity rhythm observed. To further substantiate the proposed bimodal circadian rhythm observed in BS females, a pharmacologically induced seizure model could be assessed. Wild type flies fed a proconvulsant (Stilwell *et al*., 2006) could be tested at different times of the day, thus examining the reproducibility of the bimodal rhythm in alternative seizure models.

Having established a potential influence of time of day on seizure severity of *jus* females, this study sought to ablate the central clock to determine its role in seizure rhythmicity. Under constant illumination, *Drosophila* become arrhythmic due to constant degradation of dCRY upon photoactivation (Konopka, Pittendrigh and Orr, 1989; Peschel *et al*., 2009). Hence, the molecular clock becomes arrhythmic. BS females maintained in LL conditions showed no oscillation in seizure severity, indicative of an underlying contribution from circadian physiology.

Although constant light abolished seizure rhythmicity, photic masking could be involved in *jus* seizure activity. Circadian rhythms can be “masked” by other stimuli that are presented at predictable times of day (Gall and Shuboni-Mulligan, 2022), such as light, causing the direct enhancement or inhibition of locomotor activity (Lamont and Amir, 2009). To discriminate whether seizure rhythmicity of *jus* females is circadian-mediated or masked by light, flies were reared under free-running DD conditions. Results from this suggested an effect of circadian time on seizure severity despite the absence of light stimuli, implying an effect of circadian rhythms on seizure severity of *jus* females. Studies have shown that *Drosophila* free-running circadian rhythms in DD remain stable and persist with a period close to 24 h (Plautz *et al*., 1997; Johnstone *et al*., 2022), similar to the circadian locomotor results in this study. Moreover, *jus* females maintained in free-running DD conditions exhibited a seizure severity rhythm with a longer period than in LD, thus seizure peaks were ∼4 h later in the day. Regarding the locomotor activity of *jus* females monitored in DD, this also seemed to be shifted relative to the LD rhythm. However, under DD conditions, the ‘siesta’ or inactivity period in the middle of the respective day is less pronounced (Mazzotta, Damulewicz and Cusumano, 2020). Hence, locomotor activity in DD did not show two peaks of activity like seizure severity, thus not being able to match locomotor activity to seizure severity profiles as in LD. Therefore, arguments for increased activity and nervous system excitability leading to increases in seizure severity cannot be extended to DD as in LD.

Interestingly, LL and DD conditions increased seizure severity independent of circadian time. As monitored by the TriKinetics system, such lighting regimes also led to sleep deprivation, measured by total sleep duration. Indeed, sleep deprivation is one of the most commonly reported factors associated with seizure occurrence in humans, increasing the risk of epileptic seizures (Lawn *et al*., 2014; Dell’Aquila and Soti, 2022). In addition, *jus* females in LL and DD also showed sleep fragmentation, as indicated by a reduced number of sleep episodes of shorter duration. Interrupted sleep in mice significantly increased severity of seizures (Grubač *et al*., 2021). Sleep fragmentation is considered a contributing factor to brain hyperexcitability, increasing seizure susceptibility, and offering a potential explanation to the observed increase in seizure severity of *jus* females in LL and DD. Here, it is worth noting that females used were not selected to be virgins, and mating is shown to affect sleep (Geissmann, Beckwith and Gilestro, 2019).

In humans, nocturnal light is associated with sleep disturbances and sleep deprivation, which can increase the risk of epileptic seizures (Bonilla-Jaime *et al*., 2021; Dell’Aquila and Soti, 2022). Constant light exposure also increases oxidative stress (Rodrigues *et al*., 2018), which is a critical factor involved in the initiation and progression of seizures (Shin *et al*., 2011). Sleep deprivation causes an imbalance of reactive oxygen species (ROS) production and antioxidant responses (Hill *et al*., 2018). Elevated ROS levels increase membrane excitability of *Drosophila* sleep neurones, dFB, by increasing the frequency of action potentials (Kempf *et al*., 2019). This is potentially via alterations of I_A_ currents by *Hyperkinetic*, a β-subunit of *Shaker* (*Sh*) K^+^ channels (Ueda and Wu, 2008), leading to increased neuronal hyperexcitability (Yao and Wu, 1999), and potentially increasing seizure severity. Of course, DD is an unnatural condition that may be regarded as a stressor. Mice under DD showed significant increases in stress hormones and HPA axis activation (Singh *et al*., 2023). Moreover, prolonged physical stressors such as social isolation and restraint of rodent models of epilepsy increase seizure frequency and reduce seizure threshold (Reddy, Thompson and Calderara, 2022). Therefore, a stressful environment such as DD could have increased the seizure severity of *jus* mutants.

In conclusion, this study provides evidence for an interplay between circadian rhythms and *Drosophila* seizure activity: a poorly understood phenomenon. Our findings suggest a potential bimodal circadian rhythm of seizure severity in female, but not male, *jus* files. Given that *Drosophila* is well-suited for investigating the relationship between circadian rhythms and seizure activity, future work should focus on the mechanism(s) modulating seizure rhythmicity downstream of clock genes. Excitingly, an enhanced understanding could lead to the targeting and prevention of seizure rhythmicity for the better treatment of epilepsy.

## Acknowledgements

We thank Sanjai Patel for his help with the TriKinetics system. We thank Scott Gibb and Sophie Smith for their help with Matlab for DAM analysis. We are also grateful to the members of the Baines group for their support and ideas. This work was supported by funding from a Joint Wellcome Trust investigator award to R.A.B (Grant 217099/Z/19/Z). Work on this project benefited from the Manchester Fly Facility, established through funds from the University and the Wellcome Trust (Grant 087742/Z/08/Z).

## Author contributions

Conceptualization, M.H.R.; methodology, M.H.R and A.A.B.; investigation, M.H.R.; writing – original draft, M.H.R; writing – review & editing, M.H.R., A.A.B. and R.A.B.; funding acquisition, R.A.B.; resources, R.A.B.; supervision, A.A.B. and R.A.B.

## Competing interests

The authors declare no competing interests.

## Data and resource availability

All relevant data and details of resources can be found within the article.

## Notes

### Competing Interest Statement

The authors have declared no competing interest.

